# Lipidomic Profile Reconstruction of Therapeutic Membrane Targets Using Physics-Based Optimization with Limited Activity Data

**DOI:** 10.64898/2025.12.15.694365

**Authors:** M. Krebs, H. J. Risselada

## Abstract

Membrane-targeting peptide motifs recognize characteristic features of biological membrane targets through specific interactions with their lipid composition. To elucidate the complex relationship between binding sequences and lipid composition, we propose an alternative computational framework to leverage limited sequence activity data by combining genetic algorithms with coarse-grained molecular dynamics simulations. We demonstrate through evolutionary optimization of plasma membrane lipid compositions how four true positive and four false positive antimicrobial peptide sequences encode sufficient information to resolve model membranes with a discriminative affinity of peptides that matches the accuracy of state-of-the-art antimicrobial classification. This finding reveals how subtle differences in lipid composition precisely control the selective binding of membrane-associated proteins, thereby regulating their trafficking, aggregation, and function within living cells. Our methodology reconstructs lipidomic profiles from limited sequence data, uniquely revealing targeting patterns for therapeutic peptides and the selectivity mechanisms of membrane-associated proteins.

## 1 Introduction

The interface between biomaterials such as peptide motifs and lipid membranes represents a crucial frontier in modern therapeutics, where precise molecular interactions hold the key to developing targeted treatments with minimal side effects (*1, 2, 3, 4, 5*). Beyond therapeutic applications, these interactions play fundamental roles in cellular function and protein trafficking, where membrane properties actively regulate protein organization, conformation, localization, and activity across multiple scales (*6,7,8*). This dual significance makes understanding protein-membrane interactions essential for both therapeutic innovation and basic cellular biology. Recent advances in lipidomics have revealed an unprecedented complexity in membrane composition, with over 1000 distinct lipid species in mammalian cells alone (*9*), creating both challenges and opportunities.

The significance of peptide-membrane interactions stems from their fundamental role in numerous biological processes, including antimicrobial defense, membrane fusion, protein trafficking, and therapeutic compound delivery (*10*). These interactions are particularly promising for therapeutic applications due to their ability to selectively target specific membrane types while sparing others, offering a paradigm shift in treatment strategies.

The development of effective peptide-based therapies relies heavily on our understanding of membrane composition and organization. Modern lipidomics has revolutionized this field by enabling comprehensive mapping of membrane lipidomes, allowing researchers to construct increasingly accurate model membranes for testing therapeutic candidates (*9*). This advancement is particularly crucial as membrane composition varies not only between cell types but also across different subcellular locations, necessitating precise targeting strategies (*9*).

Recent breakthroughs in peptide engineering have led to the development of highly selective therapeutic agents. The success of these approaches is evidenced by partition constants that can vary dramatically between different membrane types - for instance, certain antimicrobial peptides show significantly higher binding affinities for bacterial-like membranes compared to mammalian counterparts (*1*). This selectivity arises from fundamental compositional differences: bacterial membranes are rich in negatively charged phospholipids such as phosphatidylglycerol and cardiolipin, while mammalian membranes sequester these acidic lipids primarily in their inner leaflets (*10*). Additionally, mammalian membranes contain protective cholesterol-rich domains that confer resistance against membrane-disrupting peptides (*10*).

This emerging field of membrane selective peptides holds tremendous promise for addressing unmet medical needs through the development of targeted therapeutics with improved efficacy and reduced toxicity profiles (*11*). As our understanding of membrane lipidomics continues to evolve, we anticipate the discovery of novel therapeutic targets and the development of increasingly sophisticated peptide-based interventions that can precisely discriminate between pathological and healthy tissues (*11, 12*). While antimicrobial peptides (AMPs) have been extensively characterized with numerous well-documented sequences and mechanisms, the specific targeting of cancer cells (*13*), enveloped viruses (*14*), and clinically significant exosomes such as tumor-derived exosomes (*15*) presents a more complex scenario. This application still represents an emerging area of research, with only a small number of documented cases reported thus far (*14, 15, 16, 17*). Unlike bacterial membranes, which are highly distinctly different from mammalian cells due to their unique lipid composition based on distinct lipid types, these therapeutic targets share similar lipid types with mammalian host membranes, making selective targeting significantly more challenging (*18,19,20*).

The challenge arises from the precise lipid composition of the exposed membrane leaflet being crucial for selective peptide binding specificity and being particularly challenging to resolve (*21, 22, 20*). The limited availability of specific peptide sequences for these more difficult targets also has significant implications for the development of machine learning approaches. Traditional datadriven methods, which rely on large datasets of known examples to train predictive models, are severely constrained by the scarcity of validated targeting sequences. Several thousand active AMP sequences are currently known, yet the limited dataset size continues to challenge the development of data-driven generative models for antimicrobial peptides (*23*). This limitation highlights a critical need in the field: the development of alternative physics-based approaches that can effectively utilize the limited empirically data available to reconstruct and predict the effective lipid composition of target membranes.

This study presents a new framework that combines coarse-grained molecular dynamics simulations and genetic algorithms to optimise the lipid composition of target membranes, guided by small data. Our main interest lies in plasma membrane lipids. However, we focus specifically on antimicrobial peptides (AMPs) because they are well-studied and characterized. This allows us to validate and benchmark predictions based on extremely limited datasets against artificial intelligence-based classifications informed by large datasets comprised of thousands of labeled peptides. Such validation would be impossible in relevant domains where active sequences have only been discovered by chance, such as peptide sequences that selectively target viral membranes or tumor-derived exosomes. We demonstrate the effectiveness of our physics-based approach in leveraging minimal data to optimize a bacterial membrane model constructed from human plasma membrane lipids using a minimal dataset of four true and four false antimicrobial sequences obtained from IBM Corporation’s research on data-driven generative models for AMPs (*23*).

The effective model membrane, which has been evolutionary optimized to replicate bacterial membrane physicochemical properties while accounting for systematic force-field errors, demonstrates state-of-the-art antimicrobial classification accuracy. The versatility of plasma membrane lipids in modelling bacterial membranes highlights their key role in regulating peripheral membrane protein trafficking in cellular environments through different lipid compositions (*24*). Our concept of ‘inverse lipidomics’, which involves reconstructing effective lipid compositions for model membranes from limited activity data, shows that even a small number of membrane-selective sequences can determine membrane selectivity. Our physics-based approach, guided by minimal data, could transform the way in which sparse discoveries are integrated into the development of sophisticated peptide-based interventions that can distinguish between pathological and healthy mammalian membrane tissues with precision.

## 2 Results

### 2.1 Optimizing Lipid Compositions using Physics-Driven Evolution with Minimal Data

The primary objective of this study is to develop a physics-driven approach that optimizes mammalian lipid compositions to effectively replicate the physicochemical interactions of target membranes using minimal empirical data. With this, we aim to illustrate the discriminative power of mammalian lipid compositions in achieving selective interactions, which has significant implications for understanding the mechanisms of membrane target selectivity by peripheral membrane proteins and for designing therapeutics that can distinguish between pathological and healthy membranes in vivo.

To this end, we perform evolutionary molecular dynamics (Evo-MD) simulations (*25*), which integrate a genetic algorithm (GA) (*26*) with a parallelized coarse-grained molecular dynamics simulation framework. Unlike existing GA-based methods that often rely on neural networks and large data sets for peptide optimization (*27, 28*), Evo-MD uses a physics-based approach that incorporates MD simulations directly into the optimization process. This allows us to explore novel regions of chemical space that are not well represented in the training data, making Evo-MD particularly well-suited for scenarios with limited empirical data.

Although Evo-MD was initially designed to optimize peptides for specific membrane interactions (*25*), we adapted it to optimize lipid compositions by inverting the workflow. Using a small data set of labeled peptide sequences with known interactions, we tune mammalian lipid compositions to mimic the physicochemical properties of target membranes. This data-driven yet physics-guided approach ensures robust membrane models without requiring extensive sequence data.

For the sake of demonstration, we used a training set of eight antimicrobial peptide candidates — four true positives and four false positives — that were previously derived from a data set of 100,000 sequences generated via a deep generative model for antimicrobial discovery developed by International Business Machines (IBM) (*23*). These sequences were subsequently screened by coarse-grained molecular dynamics (MD) simulations, and the 20 highest ranked peptides were validated in vitro. Ranking was based on the effective peptide-membrane interaction measured in the simulation using a 3:1 POPC:POPG membrane model within the Martini 2 coarse grained force field (*29*). In vitro studies confirmed that two of the twenty best peptides selected in silico exhibited good activity against E. coli and S. aureus membranes, while two others showed weak to moderate activity. This example study showcases a scenario were the best 20 peptides performed similarly in the coarse-grained simulations, yet the assigned activity was only reproduced by 4 peptides in vitro. The optimization objective is to reconstruct the membrane model that would have best explained this experimental outcome using the same coarse-grained force-field.

#### Evolutionary optimization scheme

Figure 1 illustrates the basic concept of our physics-based optimization approach guided by minimal activity data. The idea is to simulate the interaction of each of the eight peptide sequences with a given target membrane and optimize the lipid composition so that the ranking based on relative membrane insertion depth aligns as much as possible with the activity ranking obtained in in vitro experiments. Similar to the original screening approach, we hypothesize that the antimicrobial activity of peptide sequences is directly reflected by their ability to insert into lipid membranes upon binding. Essentially, the degree to which a peptide inserts into a membrane reflects the tension it imposes upon binding, which is expected to correlate with its pore-forming propensity. The artificial evolution optimizes the target membrane by selecting lipids from a diverse pool of approximately 60 mammalian lipid models developed for previous simulation studies to model the human plasma membrane (*30*). Additionally, we consider optimizing differential binding with a simple model of a host membrane whose lipid composition most closely matches the average values reported in the literature for the outer leaflet of a human plasma membrane (*31*). In practice, this implies that we optimize the difference in membrane insertion depth between the host and target membrane rather than the absolute insertion depth. It is important to emphasize that the lipid composition of the host membrane remains fixed, while the lipid composition of the target membrane is the only one subject to optimization.

**Figure 1:**
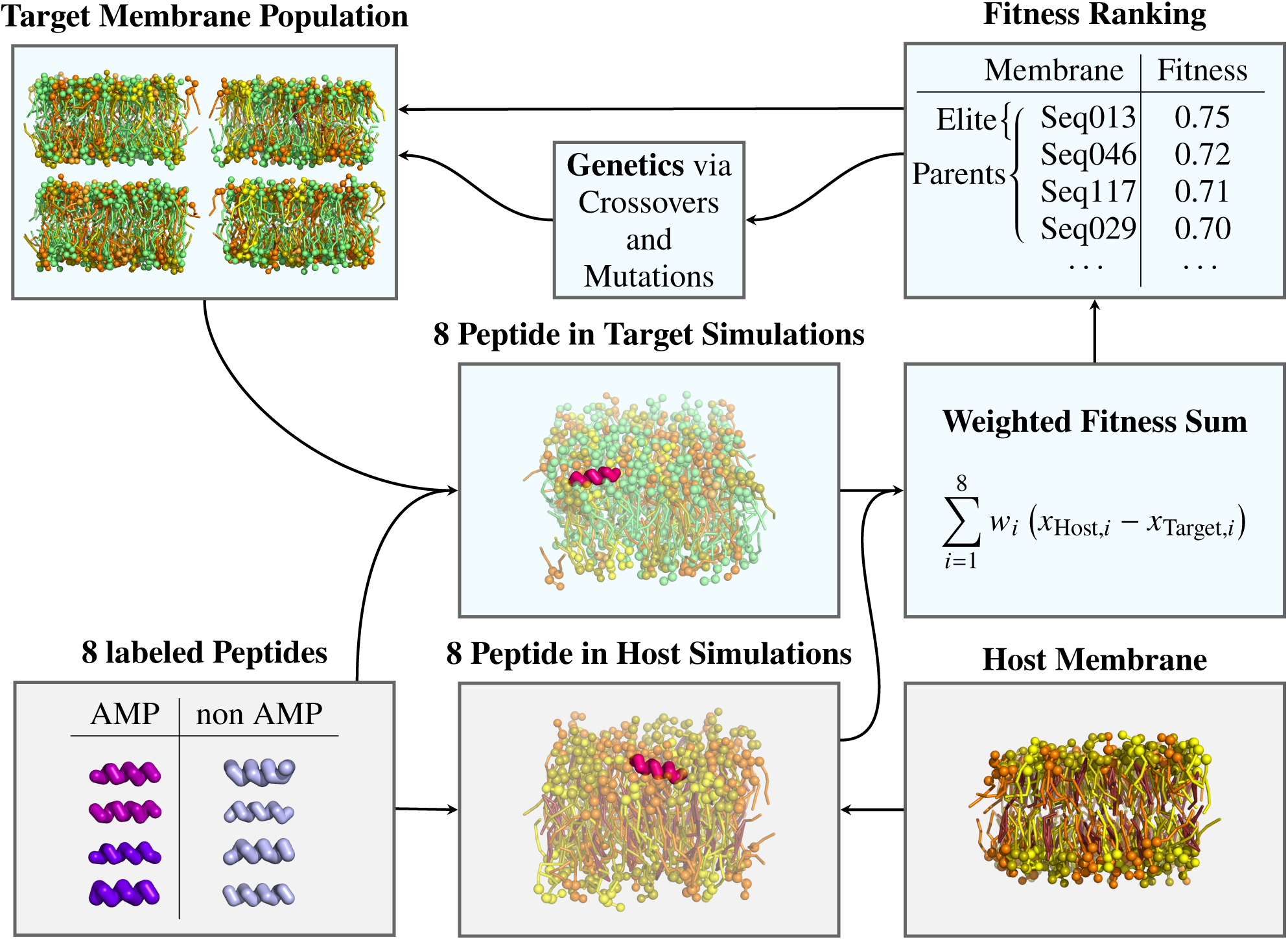
Schematic overview of the Evo-MD workflow for optimizing lipid compositions using minimal peptide activity data. Evolution starts with a random initial population of potential target membranes. The grey colored panels depict static parameters (8 labeled peptide and host membrane composition) within the optimization, obtained by averaging over three independent simulations each. The light blue colored panels (lipid composition of the target membrane) depict dynamic parameters subject to the active Evo-MD cycle. The fitness of a membrane, the relative insertion depth of Fig. 2, is calculated for each of the training peptides, via a coarse-grained MD simulation. The relative insertion depth measured for the 8 peptides are combined by a weighted sum, yielding the fitness value for the target membrane model. This procedure is carried out for all membranes of the population in parallel. The potential target membranes are then ranked by the fitness. The elite sequences are resampled directly while parents are genetically modified and their respective children form the majority of the new pool. This process is repeated, until the average fitness of the active pool converges.

### 2.2 Relative membrane insertion depth as a metric for bacterial activity

To evaluate the performance of a target membrane, we qualitatively reproduce the known ranking of eight antimicrobial peptides. The fitness function adopts maximal values when reproducing the experimental activity ranking of the eight peptide sequences (two active, two weakly active, and four inactive) in terms of relative insertion depth while maximizing the differences between them. This implies that the four active peptides maximize insertion into the target membrane while minimizing insertion into the host membrane. Conversely, the four inactive peptides (false positives) must minimize or even invert such differential insertion in order to maximize fitness (as mathematically described in section 4.3).

The empirical ranking of the 8 peptide sequences is based on the experimentally reported Minimum Inhibitory Concentration (MIC) values on E. Coli and S. Aureus (*23*). We use 8 Peptides in total, 4 of which show low MIC values and are therefore considered active, the other 4 show high MIC values and are considered inactive. These peptide candidates were originally selected from a pool of 100,000 sequences generated by a data-driven generative model. They were then specifically screened for lipid membrane interaction via in silico screening. Therefore, it is highly likely that their mechanism of action is membrane-specific. It is likely to involve either toroidal pore formation or an alternative mechanism of lipid membrane disruption facilitated by membrane insertion and leaflet tension generation.

We choose the distance *x*_target_ of the peptide to the closer leaflet of the target membrane, as a metric for describing activity in terms of differential binding affinity. For that, their positions are determined as the mean value of Gaussian fits to their respective density distributions, as can be seen in Fig. 2. Since we are specifically interested in selective binding, we also calculate the distance *x*_Host_ towards the host membrane and define the relative insertion depth as, *x* = *x*_host_ − *x*_target_ as illustrated in Fig. 2.

**Figure 2:**
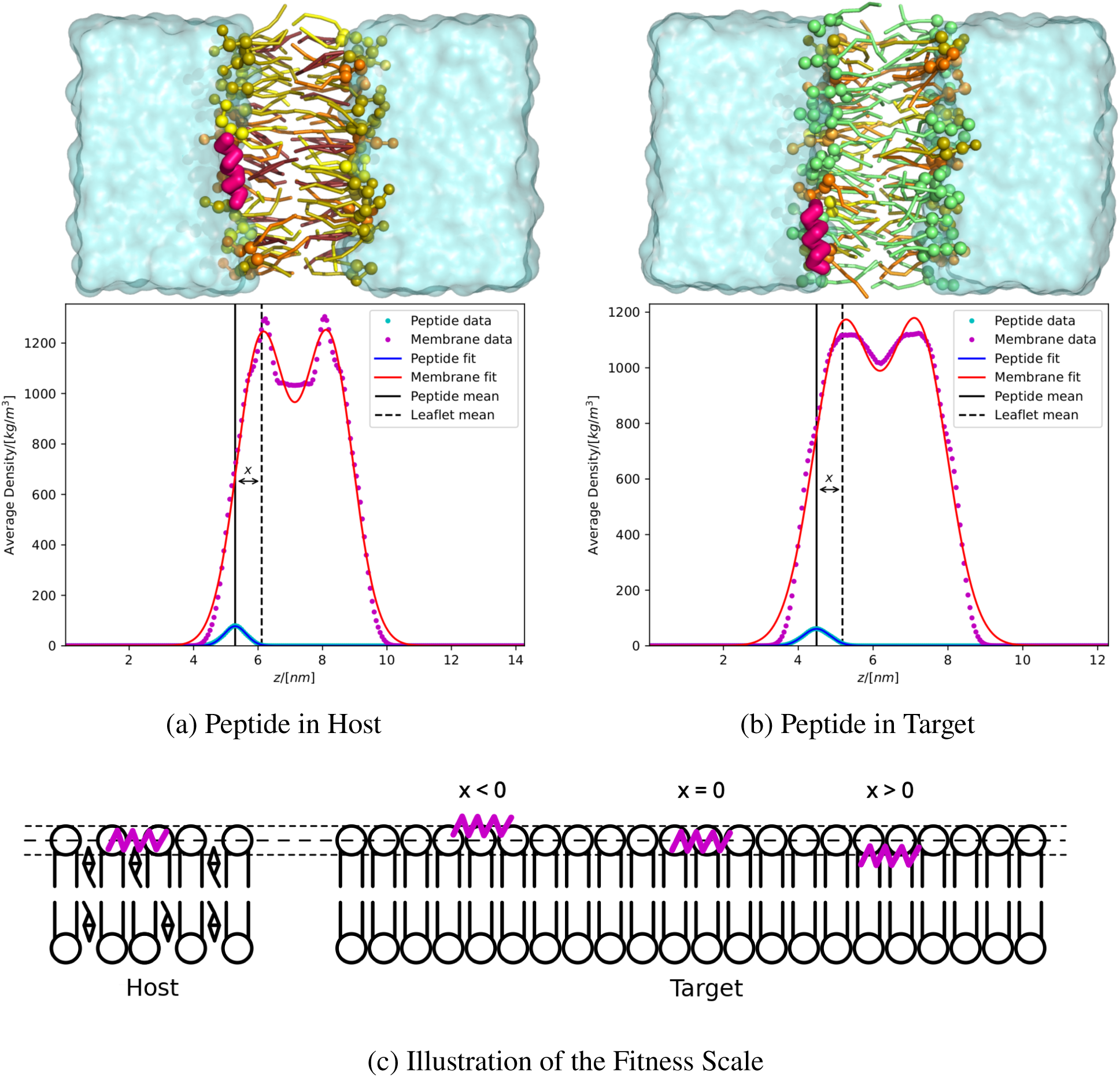
Evaluation of membrane performance. The performance of a membrane model (the fitness) is evaluated by Gaussian fits to the density of peptide and membrane. The distance of the peptide to the lipid head group plane of the membrane *x*_Membrane_ is calculated as the distance between the Gaussian mean values of the peptide (blue) and membrane (red) fits. The measured density along the z-axis is shown in magenta for the membrane and cyan for the peptide. The relative insertion depth used as a fitness to guide the evolution is defined as, *x* = *x*_host_ *x*_target_. (a) Show the fits of the peptide sequence YLRLIRYMAKMI in the host membrane. (b) Ditto, but for the target membrane. (c) Cartoon of the physical meaning of the relative insertion depth *x* for different values.

In the remainder of this work, we demonstrate that lipid compositions, when optimized using Evo-MD, can effectively mimic the discriminative interactions of bacterial membranes. This highlights the remarkable versatility of mammalian lipids in therapeutic design. The optimized membrane model achieved state-of-the-art classification accuracy for the training set, as measured by peptide insertion depths, providing quantitative insights into selectivity and potential toxicity. These findings underscore the potential of inverse lipidomics—reconstructing effective lipid compositions from minimal data—to accelerate the discovery of therapeutics capable of distinguishing pathological from healthy membranes.

### 2.3 Optimal target membrane is enriched in negatively charged lipids and depleted of cholesterol

Figure 3 illustrates how fitness improves over the course of evolution for 5 independent Evo-MD runs starting from different initial random lipid membrane compositions. The Evo-MD implementation employs an innovative asynchronous parallel processing approach, departing from traditional genetic algorithm population management methods. Rather than maintaining a fixed population size, the system utilizes an active population consisting of parallel simulations, with resource availability determining the scale of operations. In this implementation, the system leveraged 1727 parallel simulations across 1728 jobs, resulting in an effective population size of 215 lipid membrane compositions that could be processed simultaneously.

**Figure 3:**
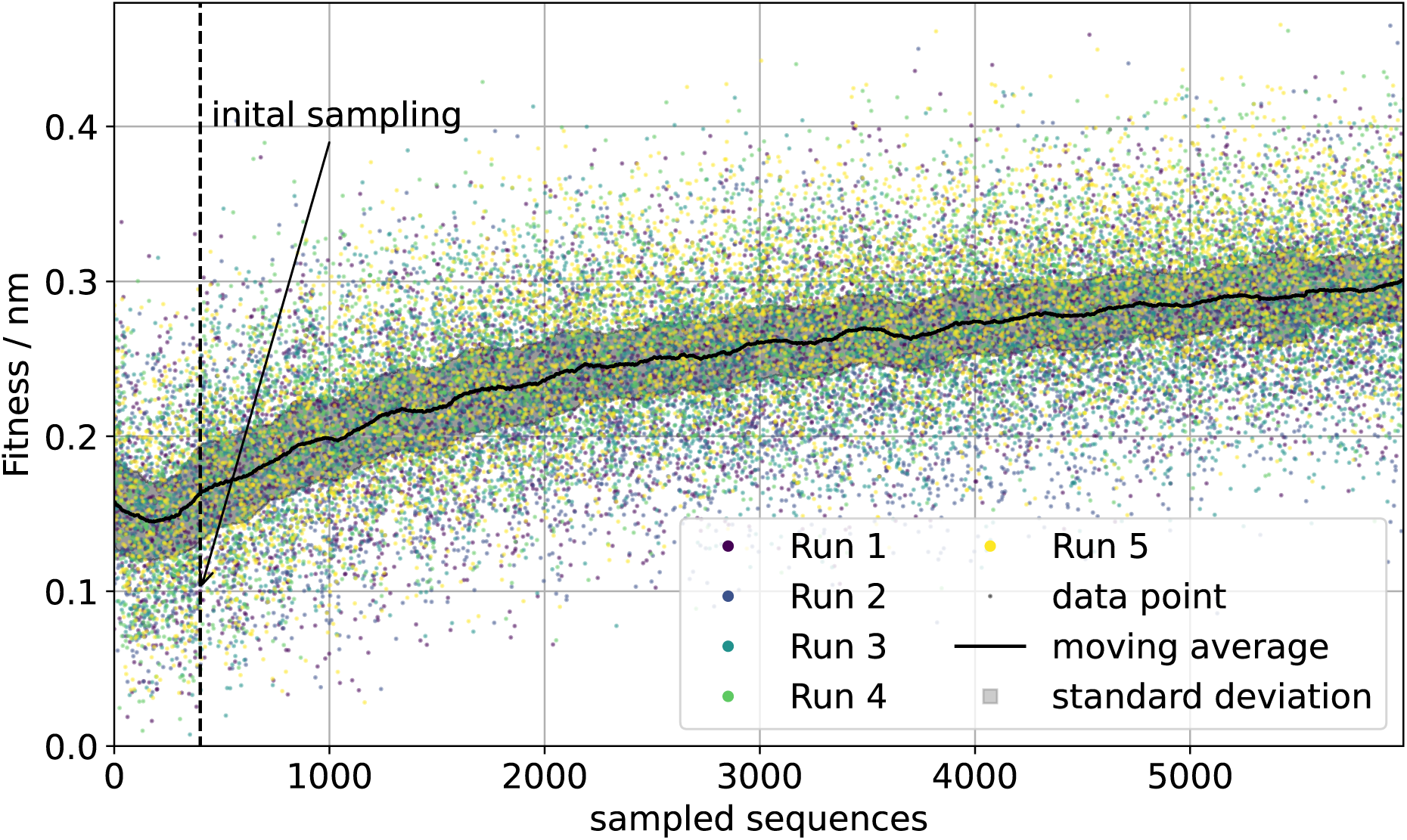
Time evolution of lipid membrane composition. The coloured dots represent the individual fitness values. for each membrane composition vector in each run, measuring the performance of the host and target membranes in discriminating the four true positive peptide sequences from the four false positive ones. The black line shows the moving average over 215 membrane composition vectors, of the five different Evo-MD runs. The grey area marks the standard deviation of this moving average. The dashed vertical line marks the end of the generation process for the initial random sampling pool, which was used to create a diverse set of membrane composition individuals and find the proper elites to drive the asynchronous genetic algorithm. Evolution stops when the composition of the overall membrane remains largely static and it is no longer justified to use computational resources for further enhancement of performance.

The best performers obtained are membrane models where the relative insertion depth of peptides shows the largest separation of true positives and false positives around a self-emerging classification cutoff (e.g., see Fig. 5). We focus on the lipid composition of “membrane 3” as a case example of a best performer (see Fig. 4a). To better clarify its lipid composition, we also plot the distributions of head group and lipid tail types separately. For reference, we also depict the lipid composition of the host membrane model in 4b.

**Figure 4:**
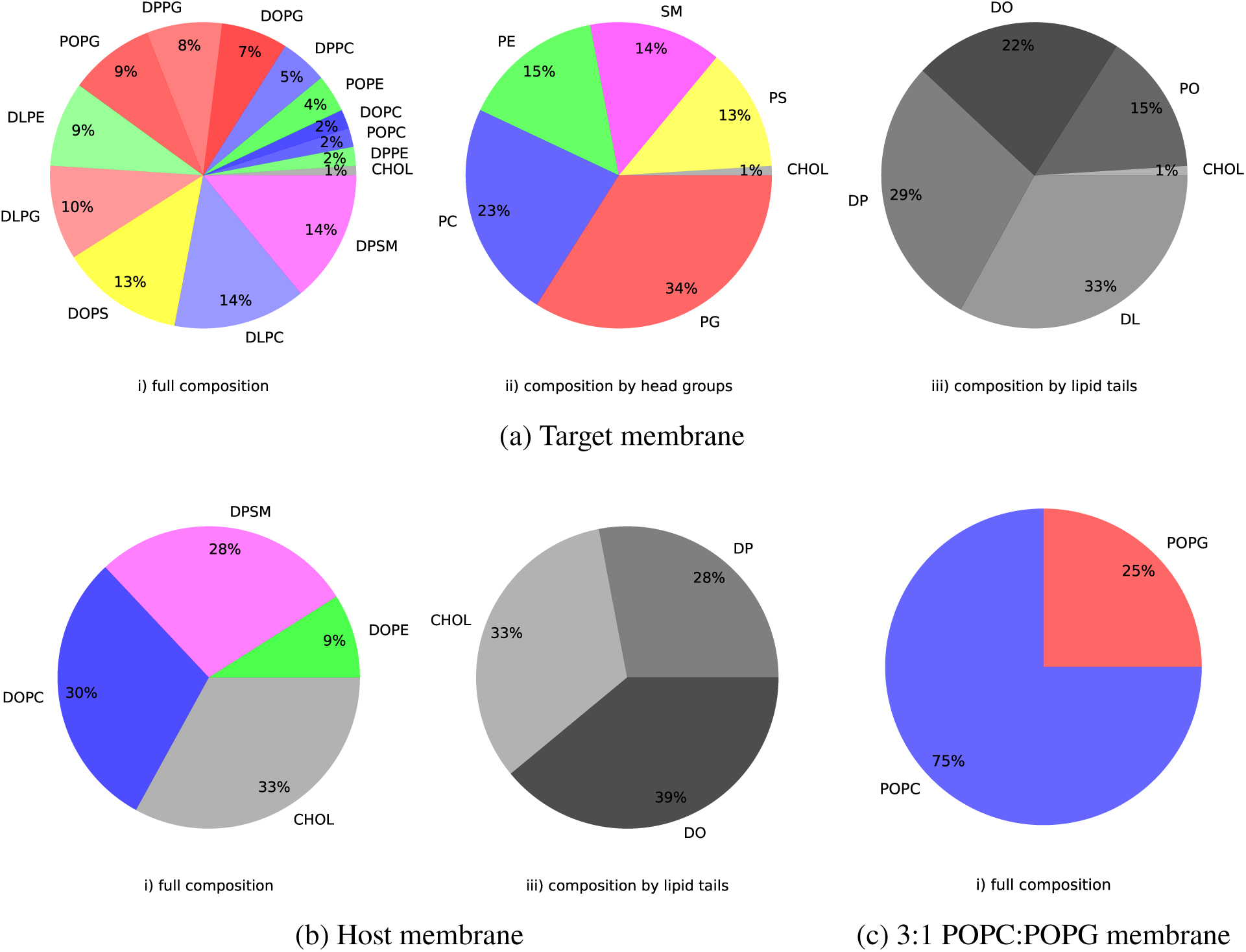
Optimized lipid membrane compositions. The Lipid distributions for the target (a), the host (b) and a stereotypical 3:1 POPC:POPG model membrane (c) membranes. The panels i) shows the complete distribution of lipids, while ii) shows the partitions of the different lipid headgroups and iii) shows the partitions of different fatty acid chains. The optimised target membrane is enriched in negatively charged phosphatidylglycerol (PG) lipids (34%) and completely devoid of cholesterol (1%), both of which contrast with the composition of the host membrane lipids. Additionally, saturated lipid tails are substantially more abundant than unsaturated ones.

**Figure 5:**
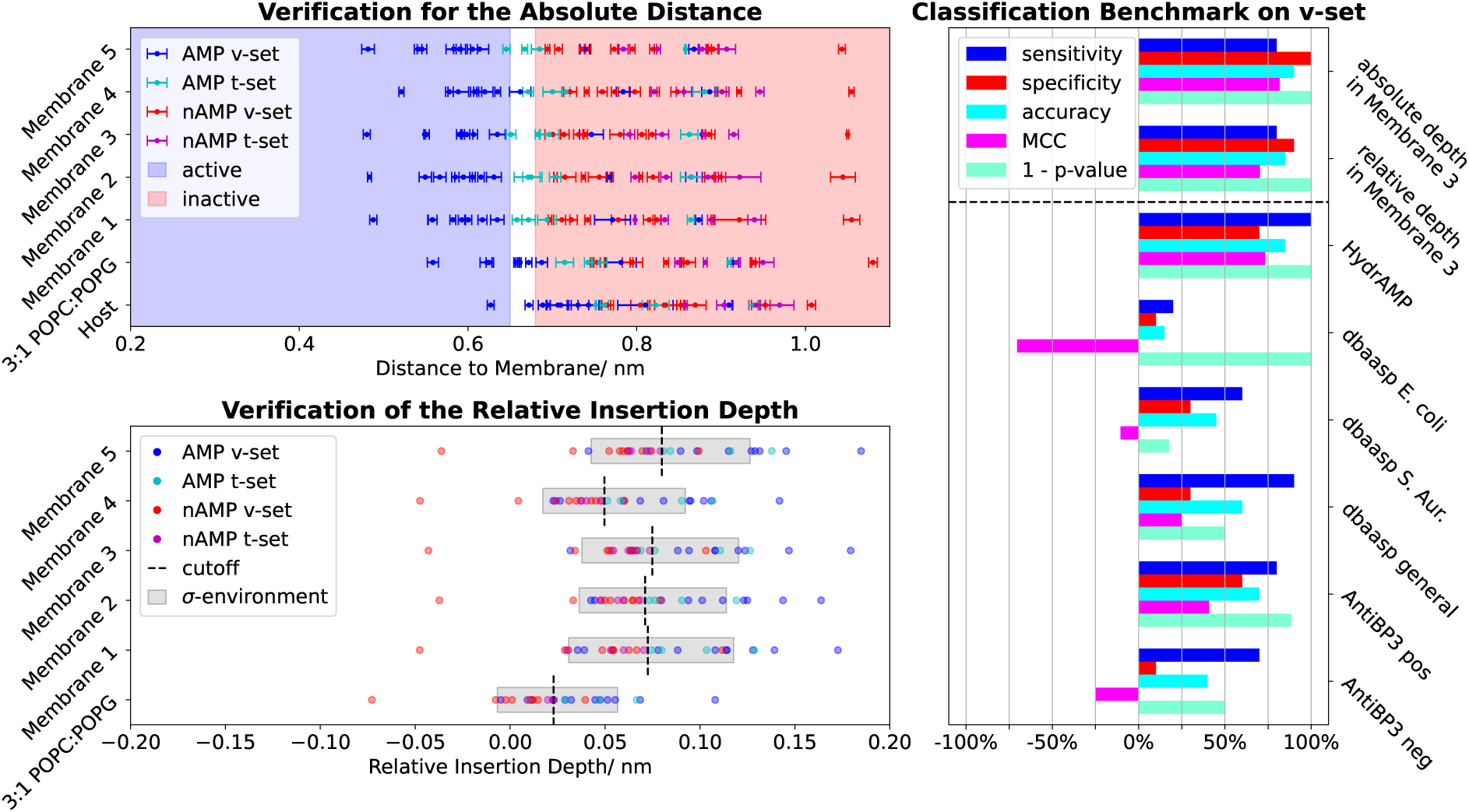
Benchmark of antimicrobial activity prediction. The performance of different membrane models was assessed by measuring their ability to distinguish between 20 independent antimicrobial peptide sequences that were empirically labelled as either false positives (red data points) or true positives (blue data points). The eight peptides of the original training set are coloured cyan (true positives) and purple (false positives). (Upper panel) *Measure of membrane binding* Plotted is the absolute distance of the peptides to the top five resulting membrane sequences, as well as the 3:1 POPC:POPG standard model and the host membrane model. The peptides are divided into those that are experimentally active (AMPs) and those that are inactive (nAMPs), and are either part of the training data (t-set) or the verification data (v-set). Each data point has been calculated three times to obtain insight into the sampling error. The blue area marks the approximate binding regime and the red area marks the non-binding regime. Values in between can be understood as indicating weak binding or an undetermined result. (Lower panel) *Measure of relative binding* The grey area indicates the standard deviation of all peptides for each membrane, showing the variation in fitness values. The black dashed line indicates the optimal cut-off point for classification within the verification data, demonstrating how this point depends on the composition. Optimised membranes demonstrate improved separation in relative insertion depth (wider grey area) compared to the 3:1 POPC:POPG membrane. (Right panel) *Benchmark of the proposed classification method, against other state of the art classifiers: HydrAMP (39), dbaasp strain specific and general classifiers (40) and ABP3 (41)* For the benchmark classification we used Membrane 3, since it shows the clearest gap on the verification set between active and not active sequences. Next to the Sensitivity, Specificity, Accuracy and Matthews correlation coefficient (MCC) we also show the p-value for the hypothesis of random classification. Our classification, as well as the HydrAMP classifier are lying in the 1% significance interval.

The predominant lipids expressed in the optimized target membrane are negatively charged phosphatidylglycerol (PG) lipids, which can be understood from the positively charged nature of the tested antimicrobial peptides (*32, 33*). Despite its similar negative charge, phosphatidylserine (PS) is expressed at much lower levels. Additional test simulations in which all PG was replaced with PS did not significantly alter the fitness values in the simulations (see Fig. S7). We therefore attribute the apparent preference for PG over PS to the fact that four times more PG species are available than PS species in the available parameter space.

Furthermore, the membranes are relatively rich in phosphatidylethanolamine (PE) and phosphatidylcholine (PC) lipids, both lipid types being commonly used in combination with PG lipids within simplified in vitro bacterial membrane models (*23*) despite PC lipids not being a common lipid type in bacteria.

Sphingomyelin (SM) and phosphatidylethanolamine (PE) are lipid headgroups that occur in smaller numbers. As we will illustrate (see Fig. 4c) selectivity can largely be obtained by a membrane consisting solely of two majority components, phosphatidylcholine (PC) and phosphatidylglycerol (PG), given that the fitness is based on the differential binding with the host membrane. Therefore, the contribution of lipids like SM and PE to antimicrobial selectivity does not seem crucial. Finally, the appearance of cholesterol is small enough to be neglected within the statistical boundaries of genetic operations, yielding an effectively cholesterol-free membrane, as one would expect of a bacterial membrane. This strongly contrasts with the 33% cholesterol present in the host membrane model (see Fig. 4b). Cholesterol thus plays a major role in the selectivity of antimicrobial peptides. Its well known effect on increasing the ordering of lipid tails in eukaryotic membranes illustrates this role. This impairs the insertion of antimicrobial peptides into a eukaryotic membrane, resulting in an active protection against the membrane lytic action of antimicrobial peptides.

Concerning the expression of lipid tail types in the optimized target membrane, the most abundant tail types are the saturated DL (diacylglycerol lipids) with 33% and the longer DP (diacyl phosphatidylcholine) lipid tails with 29% forming the majority constituents of the target membrane. The strong expression of saturated lipid tails substantially differs from eukaryotic membranes that largely consists of mono (PO) and double unsaturated (DO) lipid tails. This expression is highly consistent with the notion of many bacterial target membranes such as Staphylococcus aureus (S. aureus, Gram-positive) being almost entirely consisting of saturated lipids (*34*) whereas Escherichia coli (E. coli, Gram-negative) features a more balanced mix consisting of 50-60 % saturated lipids (*35, 36*). This however still substantially contrasts the human plasma membrane which only contains up to about 20-25 % saturated lipids. The prevalence of saturated lipids in the bacterial membranes likely compensates for the absence of cholesterol and the prevalence of non-lamellar phase forming lipid types such as lysolipids and PE lipids by providing mechanical stability to the membrane. At first glance, it may seem surprising that evolution favors saturated lipids, given that unsaturated lipids intuitively produce softer, less stable membranes that are more susceptible to peptide action. However, saturated lipids increase the hydrophobicity of the membrane surface by creating more defects in the packing of hydrophobic lipid tails at the oil-water interface (*37*), which consequently reduces hydration repulsion (*38*). In contrast, unsaturated lipids mitigate these defects, consequently reducing surface hydrophobicity (*38, 37*). This increased surface hydrophobicity strengthens the interaction between amphiphilic peptide sequences and the membrane surface, making it thus more susceptible to destabilization by amphiphilic surface-active peptides.

Our results provide a first striking example of how limited data on the activity of eight sequences can be used in conjunction with information on protein-membrane interactions encoded within (coarse-grained) bio-molecular force-fields to reconstruct the lipidome profile of an effective target membrane. The obtained profile contains features that can be intuitively linked to the known properties and characteristics of bacterial membranes. Our results demonstrate that scarce data on activity is sufficient to reveal how selective peptide sequences and membrane binding motifs “visualize” differences between biological lipid membranes, the basis of targeting.

### 2.4 Optimized membrane compositions leverage antimicrobial peptide classification accuracy

Analysis of the solution space of the optimal target membrane (Fig. 4a) reveals that a minimal bacterial membrane model primarily contains two components: phosphatidylglycerol (PG) and phosphatidylcholine (PC). This is consistent with the composition of typical in vitro screening assay membranes. However, all of the peptides used in the training set originate from a study that classified them as true positives based on in silico screening using a 3:1 PC:PG membrane (*23*). We attribute this distinction to the different metrics used to determine antimicrobial activity in the referenced study. In the referenced study, the metric was contact between positively charged peptide residues and negatively charged lipid head groups (*23*). In our study, the metric is membrane insertion depth.

To further verify the classification capabilities of the resolved membranes, we tested a set of 20 independent, labeled antimicrobial peptide candidates that were not included in the original small activity dataset used to optimize the membrane. These peptides consisted of 10 known active AMPs (true positives) and 10 false positives generated by the IBM deep learning model (*23*). The 10 active AMPs were randomly chosen from the APD3 antibacterial database (*42*) under a few conditions. They had to be active against either E. Coli or S. Aureus and conform with the in vitro screening performed in Ref. (*23*). Additionally, they had to have a helical secondary structure and be shorter then 20 amino acids to prevent the occurrence of system size effects on membrane insertion within the small simulation setup. The 10 false positives were predicted as true positives in the in silico screening and were then tested experimentally. Therefore, they serve as hard targets in our validation approach. As a point of reference, the same classification was performed on the original 3:1 POPC:POPG membrane that was used for in silico screening.

In Fig. 5 (upper panel), the membrane affinity as characterised by the membrane insertion of all 20 peptides into the five best membranes, optimised by Evo-MD, are shown. The 3:1 POPC:POPG membrane and the host membrane are shown for reference. Active peptides are plotted as red data points and inactive as blue data points. We define two regimes matching with the experimental labels. We mark the high-affinity regime with a blue shaded area and the low-affinity regime with a red shaded area. There is a small boundary area in between where the classification is undetermined.

For the AMP-labelled set from the APD3 database, it is clear that most of the active peptide sequences fall in the high-affinity regime in the top five Evo-MD membranes. For the negatively labeled peptides it can be seen, that they generally fall in the low affinity regime, as one would expect. However, for the 3:1 POPC:POPG membrane, they shift slightly towards the boundary regime, indicating weaker preferential binding interactions than with Evo-MD-trained membranes. One active sequence falls within the strong affinity regime for the host membrane suggesting the potential for toxicity. This peptide is the natural version of XT-7, which indeed exhibits significant haemolytic activity in cell experiments (*43*). Notably, the positively labeled training peptides used to optimize the membrane compositions show only weak to moderate membrane affinity. This is in line of expectation, however, since these AI generated sequences only showed weak to moderate activity in experiments with MIC values down to 31 µg mL^−1^ for E. Coli and 7.8 µg mL^−1^ for S. Aureus (*23*). It must be noted that the internal ranking of the four positively labeled training data, sorted by their MIC value, is not reflected by their membrane affinity in simulations.

Next, we assess the relative insertion depth into individual membranes compared to the host membrane. Active peptides generally fall on the left side of the plot (deeper insertion in the target membrane than in the host membrane), whereas inactive ones fall on the right. All of the tested membranes enabled the selection of a cutoff point for the binary classification of activity, with at least 17 out of 20 peptides being correctly classified. While the binary classification performance of all membranes was similar, the location of the cut-off point proved to be highly dependent on the membrane model. This illustrates the importance of activity calibration in modelling membranes. Interestingly, even the original 3:1 POPC:POPG membrane showed a similar binary classification performance to the five optimised membranes. However, its reduced affinity for the membrane results in a much smaller spread between data points, making classification more prone to errors. This reduction in discriminatory ability is reflected by a smaller spread in relative insertion depth, as indicated by the shaded grey area in the lower panel of figure 5.

Two important conclusions can be drawn from the observations: First, selectivity is largely explained by the two majority lipid components (phosphatidylcholine (PC) and phosphatidylglycerol (PG)) and the absence of cholesterol. Second, though the absolute depth of peptide insertions and the spread herein differ between tested membranes, the ranking of peptide insertions is mostly conserved.

The main outcome of the fitness optimization is an increased spread in relative membrane insertion/affinity (see Fig. 5), as a narrower spread reduces discriminative capacity by making predictions more susceptible to statistical flukes and force-field inaccuracies. Importantly, our results showcase that this enhanced discriminative capacity, derived from just a few peptide sequences, carries over to other independent antimicrobial peptide datasets. Our findings further illustrate that the PC:PG bacterial membrane models commonly used in in silico and biophysical screening assays, despite their simplicity, in principle provides a reasonable model for the action of antimicrobial peptides. In addition, including the host membranes in the screening process makes it easier to distinguish between peptide activity (high affinity for the target membrane only) and peptide toxicity (high affinity for both membranes), which is crucial for the optimization of therapeutic peptides.

Finally, performance was compared against state-of-the-art machine learning approaches: HydrAMP (*39*), dbaasp (*40*) and AntiBP3 (*41*) (Fig. 5, right panel). We defined five key metrics (section 4.4): sensitivity (true positive prediction rate), specificity (false positive rejection rate), accuracy (overall prediction quality), Matthews correlation coefficient (A common metric for (machine learning) classifiers (*44, 45*)), and p-value (randomness indicator). Our method achieved the highest scores in accuracy as well as specificity due to correct false positive predictions, the area were our physics-based approach excelled. Notably, the training set’s false positives were generated by a competitive data-driven generative model (*23*), making them particularly challenging for other data-based classification methods. Furthermore, our method achieved second-highest sensitivity, closely following HydrAMP, which correctly identified all true positives. Despite being optimized with only 4 active and 4 inactive sequences, our physics-based classifier thus performed comparably to state-of-the-art data-informed methods trained on thousands of active antimicrobial peptide sequences. This demonstrates that limited activity data in new, unexplored domains involving selective protein-membrane interactions can successfully achieve high classification accuracy when enhanced with physical principles. The model’s ability to accurately screen false positives represents its most valuable quality, as this significantly streamlines experimental workflows by eliminating unnecessary testing procedures. Traditionally, false positives results are viewed as failures or noise in the experimental process. However, our approach leverages the fact that these false positives can contain valuable information that improves future predictions. Furthermore, the quantitative predictions of the distances are expected to correlate with activity and toxicity. This metric can be exploited by genetic optimization towards optimal sequences via high-throughput simulations, as demonstrated in our recent works (*46, 25*).

## 3 Discussion

Here, we utilized physics-based optimization of lipid membrane composition, guided by small-scale activity data. Our methodology demonstrates how a restricted dataset of known active and inactive sequences, combined with biomolecular force fields and genetic algorithms, yields distinctive insights into membrane binding sequence recognition patterns.

Peripheral membrane proteins often contain specialized structural motifs that enable specific recognition and interaction with distinct lipid compositions in cellular membranes (*47, 48, 49*). These motifs fulfill essential roles in protein targeting, cellular signaling, and membrane trafficking processes (*50, 51*). Our methodology enables detailed reconstructions of lipid compositional patterns targeted by membrane-binding motifs. This includes the membrane-directed binding mechanisms of intrinsically disordered proteins and their subsequent membrane-controlled nucleation into protein condensates (*52,53*). We therefore expect that the here-presented concept can strengthen the understanding of the intricate relationships between amino acid sequences and membrane binding specificity.

Substantial evidence indicates that the outer leaflets of cancer cells and enveloped viruses likely exhibit higher expression of negatively charged lipid species, elevated cholesterol levels, and increased PE lipid expression (*54, 55*). However, more subtle insights into the characteristics of the outer leaflet that are essential for targeting distinct cancer cells and enveloped viruses are not readily accessible through experimental approaches. In addition, even at the overall membrane level, lipid compositions have been shown to vary greatly between different strains of influenza viruses (*20*). Our approach focuses on exploiting rarely discovered sequences with active broad-spectrum activity to effectively overcome the uncertainty and variability of leaflet lipid expression reported in lipidomics studies. This allows us to develop enhanced models that improve both biophysical and computational screening methodologies.

Furthermore, our empirically guided optimization approach offers a crucial advantage by addressing systematic biases inherent in biomolecular force fields, particularly in coarse-grained models like the Martini force-field. These biases can either overestimate or understate protein hydrophobicity, but a data guided approach effectively mitigates these issues by effectively incorporating them into the optimized lipid composition, thereby minimizing their impact on peptide design. This bias correction mechanism has significant implications for activity prediction of antimicrobial sequences, as it explains why differential binding between target and host membranes proves more predictive than target membrane affinity alone. This property becomes particularly valuable for therapeutic applications such as targeting cancer cells or enveloped viruses, where simplified models of human lipid types and differential compositions of target and host membrane must be adjusted to enable selective interactions while accounting for systematic force-field errors and model inaccuracies.

The performed benchmarks revealed that optimized membrane configurations significantly enhanced the spread in fitness across independent datasets, resulting in more robust classification capabilities. While the inclusion of bacteria-specific lipids (such as cardiolipin) could yield additional qualitative or quantitative improvements (*56, 57*), our findings demonstrate that a basic set of mammalian lipids already enables effective peptide sequence classification. This discovery reveals an interesting principle: the mammalian base lipid set can effectively access the bacterial membrane subspace through its structural and physiochemical properties as an ensemble, rather than through exact compositional matching.

The fact that the set of mammalian base lipids reproduces bacterial membrane properties so well suggests that membrane selectivity in mammalian cell membranes can be encoded in many different ways, and that interaction with unique membrane-binding motifs can be made highly selective. It further supports the possibility of selectively targeting viral, fungal, and cancer membranes. The mammalian lipid set demonstrates remarkable versatility in modeling various structural and physicochemical properties. This observation challenges the prevailing view that precise modeling and duplication of target membrane specificity via target-specific lipid types is essential. Instead, it suggests shifting the focus of predictive membrane models for therapeutic peptide screening toward exploiting experimentally labeled peptides.

The current study only used data on the labelled activity of eight sequences. This implies that eight simulations were performed for each membrane composition. In order to efficiently incorporate extensive data sets on labelled peptide sequences without escalating computational expenses, we propose the implementation of random selection schemes, which would facilitate the continuous selection of peptides from an expanded pool of labelled target peptides. This approach allows for an extensive dataset of experimentally labeled peptides during optimization, extending beyond just a few peptides.

Finally, our approach demonstrates significant potential for designing membrane-targeting therapeutic peptides using optimized target membrane models in physics-based sequence design (*25, 46*). By combining physics-based membrane composition optimization guided by small experimental data with evolutionary molecular dynamics of peptide sequences (*25, 46*), we establish a robust and unique framework for improved rational therapeutic peptide design. This methodology offers promising solutions for addressing antimicrobial resistance and viral epidemics while maintaining precise control over peptide properties and membrane interactions. In particular, the potential toxicity or activity of peptides can be inferred quantitatively from their interactions with host and target membranes. Thus, the evolution of peptide sequences can be directly tuned to optimise these properties by maximising target membrane affinity and minimising host membrane affinity simultaneously. In addition, sequences generated during genetic optimization can be pre-screened using existing data-informed classification methods (*39*) to bias sequence pool generation toward peptides with high antimicrobial propensity. In turn, similarly to our recently released PMIpred server (*58*), which uses an transformer architecture informed by large evo-MD data to classify membrane nonbinding, membrane curvature sensing and general membrane binding for user-provided peptide sequences, our approach can enhance state-of-the-art, data-informed classification tools by providing an independent, strong, orthogonal classifier that excels at predicting false positives.

## 4 Materials and Methods

### 4.1 Asynchronous Evo-MD

Evo-MD is a genetic algorithm (*26*) which obtains its fitness values by carrying out coarse grained MD simulations. It is implemented in python 3 (*59*) and parallelized by mpi4py (*60*) and OpenMPI (*61*). It was first used in a synchronized version, where the genetics were carried out in the beginning of each generation (*25*) To mitigate downtime of compute nodes, stuck idle and waiting for the rest of a generation’s population to finish, we implemented an asynchronous framework for our genetic algorithm. Instead of having one new population each iteration, there is one united sequence pool, of the best performing sequences so far, serving as parents. We start of with an initial sampling of 400 random sequences. For the membrane optimization we chose a size of *N* = 20 for this pool. Whenever a sequence finishes, the pool is updated and a new sequence for the finished process is chosen on the fly. The elite sequences, which in this case are the top 5 sequences in the pool, are rerun up to a total of 3 runs to validate their results. The rest of the parent sequences is rerun up to two times.

### 4.2 MD-Framework

The MD simulations in this study were carried out in the coarse grained (CG) force field Martini 2 (*29*) in the simulation engine Gromacs (*62*). Using a coarse grained parametrization is crucial for doing high throughput screening in a reasonable amount of time. This need for coarse graining is one of the reasons why the specific training set of (*23*) has been chosen. Vice versa the choice of the training peptides is the reason, we use martini 2 instead of the newer martini 3 (*63*), since we want to keep the exact same parametrization.

The initial simulation box is built by ABSOLUTE INSANE, a slightly adapted version of INSANE (*64*), which allows absolute numbers for lipid types instead of ratios. It contains a membrane of 100 lipids per leaflet as well as martini water with a 0.15 mol/l salt concentration.

The peptides are build separately by the python module PeptideBuilder (*65*), reparameterized by martinize2 (*66*) and aligned by their axes of inertia using the principal axis theorem (*67*) implemented via numpy (*68*).

By using a lipid map, created by ABSOLUTE INSANE, some central lipids of the membrane are pushed aside, while assuring no overlap of any lipids. This creates an empty space, where the peptides can be inserted parallel to the membrane surface.

The simulation cycle starts with an Energy Minimization (EM), Equilibration (EQ) loop in the NPT ensemble, with a timestep of 1 fs, using the berendsen thermostat and barostat (*69*), to close all the molecular cavities within the grid-like structure resulting from the INSANE generation of the system. This cycle is ended on an EM followed by a pre-equilibration with 20fs time steps for 50ns using the berendsen thermostat and barostat. After the system has stabilized, the thermostat and barostat are replaced by the more accurate v-rescale thermostat (*62*) and the Parinello-Raman barostat (*70*). The system is then equilibrated for another 50 ns, enabling additional time for the membrane mixture to reach stability conditions with the new thermostat and barostat. Finally the actual production run of 900 ns is being carried out and the densities of the peptide and the membrane lipids are calculated via the Gromacs density tool (*62*). We chose 900 ns of production after testing different runtimes and analyzing the convergence of the fitness, which can be seen in the supplementary Figure S8.

### 4.3 Fitness evaluation

To calculate the peptide-membrane distance, we measure the density in z-direction (orthogonal to the membrane surface) over the whole production run using the gromacs density tool (*62*). We use Gaussian least square fits

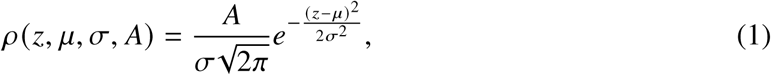

carried out via scipy’s (*71*) curve_fit function, to determine the average position of the peptide *μ*_p_ and both membrane leaflets *μ*_l1_ and *μ*_l2_, as can be seen in figure 2. We define *x* as the distance between the peptide position *μ*_p_ and the closest leaflet position *μ*_l_. The sign of *x* is positive for the peptide outside the membrane and negative for the peptide inside the membrane.

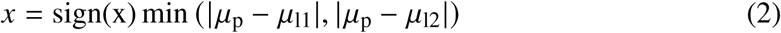

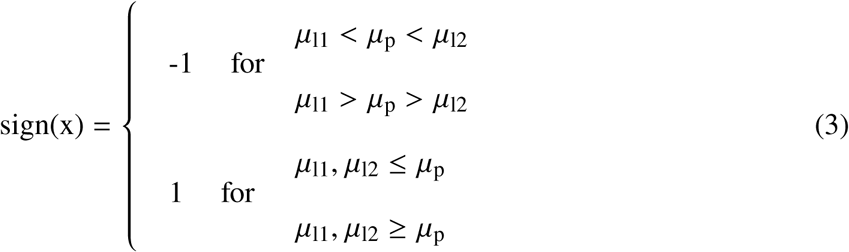

For all 8 peptides we compute their distance *x*_target_ to the bacterial membrane model in question and their distance *x*_host_ to the human membrane model of (*31*). From that we define the relative insertion depth *x*_rel_ = *x*_host_ − *x*_target_ The fitness *F* of a bacterial membrane is now a weighted sum over the relative insertion depth of all 8 peptides into the membrane

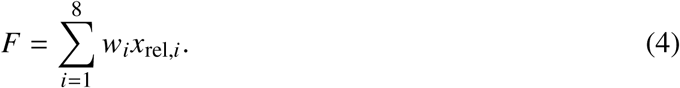

The weights *w_i_* are chosen to be positive for experimentally active peptides (true positives) and negative for experimentally inactive peptides (false positives). Therefore, a higher fitness means a better agreement with the qualitative ranking of the peptides. To prevent fitness optimization by general peptide attraction or repulsion instead of selective behaviour, we enforce the constraint

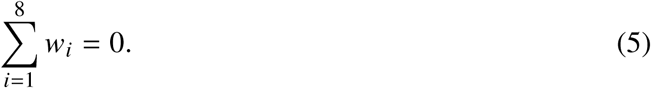

### 4.4 Statistical classification metrics

For verifying and benchmarking our method, we applied multiple different classification metrics. For this, we divide the verification data into four different groups:

TP: true positives, the number of correctly identified active peptides. TN: true negatives, the number of correctly identified inactive peptides.

FP: false positives, the number of peptides that were classified as active, but really are inactive. FN: false negatives, the number of peptides that were classified as inactive, but really are active.

Firstly we define the sensitivity (*72*)

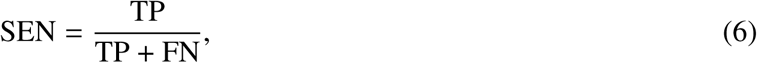

which measures how well a classifier performs on truly active peptides; The specificity (*72*)

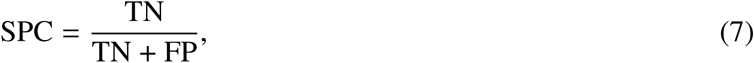

which measures how well a classifier performs on truly inactive peptides; And accuracy (*72*)

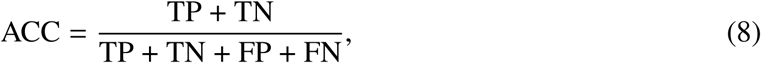

which measures the rate of true predictions. These three metrics result in values from 0 to 1, where 1 indicates perfect predictive behaviour, 0.5 shows random behaviour and 0 means perfectly anticorrelated behaviour. For the accuracy metric to be relevant, it must be noted, that the verification set needs to be balanced, meaning it needs as many truly positive peptides as it has truly negative ones. We satisfy this requirement by choosing 10 active and 10 inactive peptides for our verification.

Secondly, we define the Matthews Correlation Coefficient

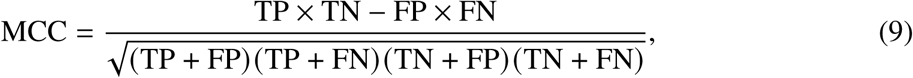

which is another metric for the overall prediction quality of a classifier. The MCC factors in all basic rates of the binary classification and only yields a high value if all of them are performing sufficiently well. Therefore, it is used as the gold standard for (machine learning) classifiers (*44,45*). The MCC reaches from -1 to 1, where 1 indicates perfect prediction, 0 means random behavior and -1 shows perfect anti-correlation.

Lastly, we apply a two-tailed p-value test (*73*) for the hypothesis that our results are random. We want to show that it is very likely that the classification on a comparatively small verification data set is not only successful because of a lucky pick of verification peptides. The two-tailed p-value for *x* correct classifications within *N* tries is calculated as a sum of the probabilities of all possible outcomes in both tails.

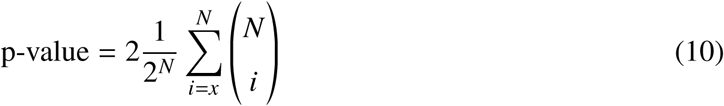

for *x > N*/2, or

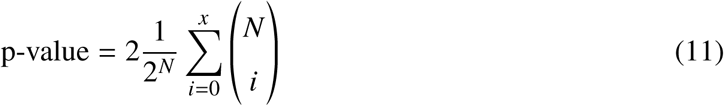

otherwise. The binomial coefficient is defined as

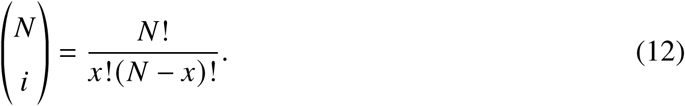

The hypothesis can then be rejected, if the p-value lies under a certain threshold which in our case is 0.01. While the exact implication of the p-value test is under discussion (*74*), the rejection of our random results hypothesis should at least be an indicator for our positive classification results to not be results of random lucky picks.

## Acknowledgments

The authors gratefully acknowledge the Gauss Centre for Supercomputing e.V. (www.gauss-centre.eu) for funding this project by providing computing time through the John von Neumann Institute for Computing (NIC) on the GCS Supercomputer JUWELS at Jülich Supercomputing Centre (JSC).

The authors thank Dr. Jeroen Methorst and Sebastian Lütge for their contributions in the Development of Evo-MD.

## Author contributions

H.J.R. designed and supervised the project. M.K performed all simulations and analysis. Both authors wrote the manuscript.

## Data and materials availability

Data and developed source codes will be uploaded on a publicly accessible Github.

## Supplementary materials

Figs. S1 to S8

Tables S1 to S12

