## Supplementary Information for "Lipidomic Profile Reconstruction of Therapeutic Membrane Targets Using Physics-Based Optimization with Limited Activity Data"

M. Krebs, H. J. Risselada*

**This PDF file includes:**

Supplementary Text

Figures S1 to S8

Tables S1 to S12

### 5 Supplementary Text

#### 5.1 Convergence

To demonstrate the convergence of the algorithm, we show the development of reruns of the top-performing sequences from one run, compared to the slope of fitness development throughout one run. As shown in Fig. S1, the variation of the fitness value of a single sequence is significantly higher than the increase in fitness after sampling 5000 sequences. Therefore, we don't expect a significant gain in a reasonable amount of time. In that sense, we consider this run to be converged.

#### 5.2 Verification Data

The following plots, Fig. S2 to S6, show the composition of the top 5 resulting membranes The following tables, Table S1 to S12, show the relative insertion depth $\Delta x$ as well as the peptide-membrane distance $x$ and the peptide-host-membrane distance $x_{\mathrm{Host}}$ for all training and verification peptides for the top 5 resulting membrane compositions as well as the standard 3:1 POPC:POPG model membrane. All values are averaged over three independent simulations. The optimal cutoff for the relative insertion depth was chosen as the midpoint between neighboring peptide values that offered the most correct classifications. Upright digits mark right classification, italic digits mark wrong classification. The cutoff for activity in absolute terms, has been empirically chosen as 0.7nm. Values that indicate activity are marked by a $*$, values that indicate toxicity are marked by a †.

#### 5.3 PG and PS lipids

To illustrate the difference between the PG and PS lipid headgroups, we replaced all the PS lipids in Membrane 3 (Fig. S4) with PG lipids, and vice versa. We then calculated the fitness of these two membranes multiple times and compared them, as shown in Fig. S7. The small difference between the two membranes indicates that the PS headgroup may be slightly preferred. However, the difference between these membranes is not significant. Therefore, we conclude that our model cannot properly resolve the difference between these headgroups at the coarse-grained level.

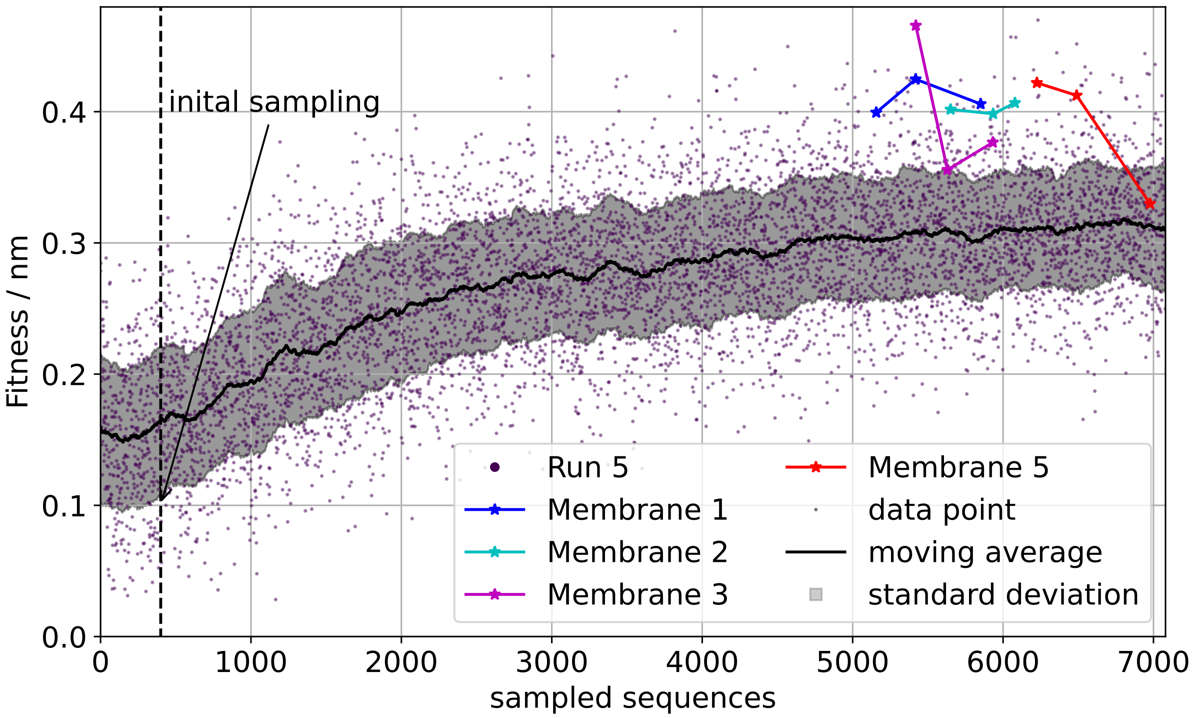


**Figure S1**: Convergence of fitness of ”run 5” is shown. This run produced the best individuals overall, including Membranes 1, 2, 3, and 5. Their occurrences in the Evo-MD history of Run 5 are highlighted. Particularly, the fitness variation of membrane 3 and membrane 5 reruns is high compared to the average fitness development at that time. There is also no gain in fitness from membrane 1 to any of the other membranes, despite the fact, that membrane 1 is the first ranked membrane to occur.

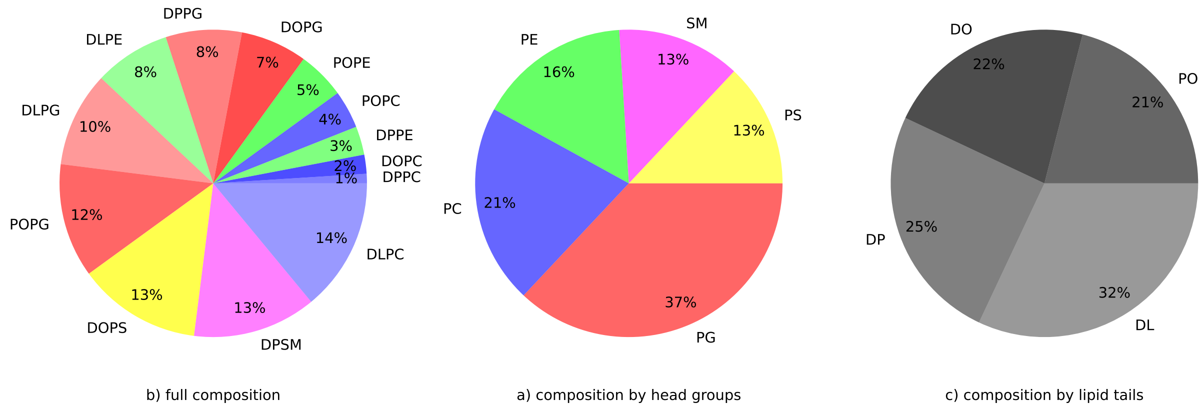


**Figure S2**: Membrane 1 Composition.

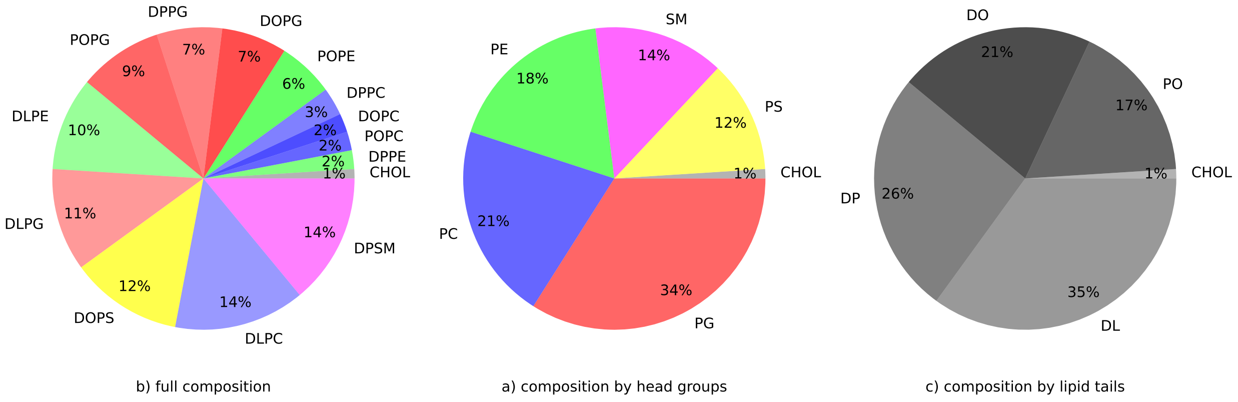


**Figure S3**: Membrane 2 Composition.

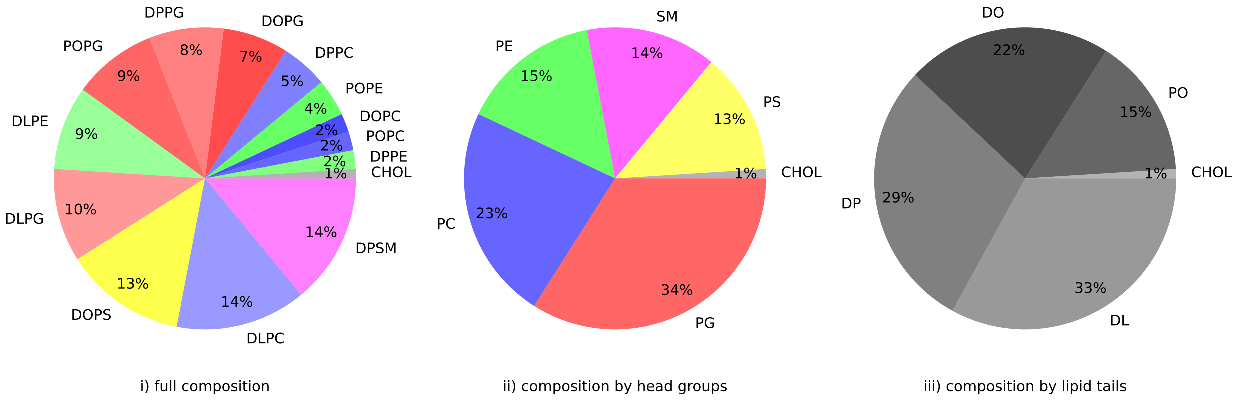


**Figure S4**: Membrane 3 Composition.

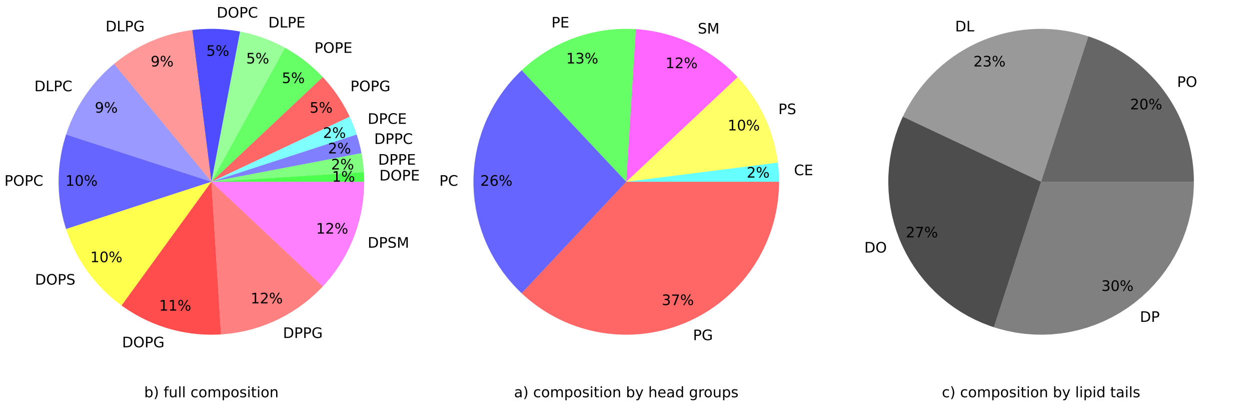


**Figure S5**: Membrane 4 Composition.

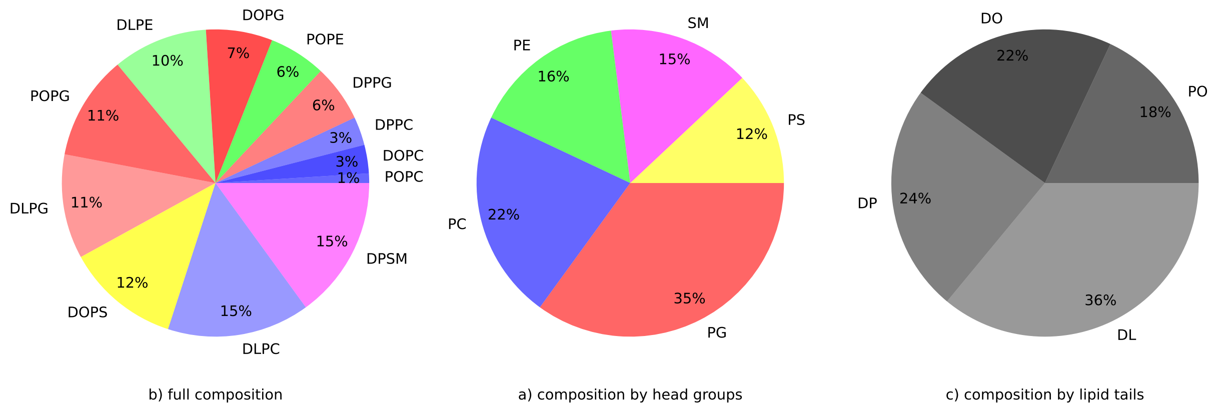


**Figure S6**: Membrane 5 Composition.

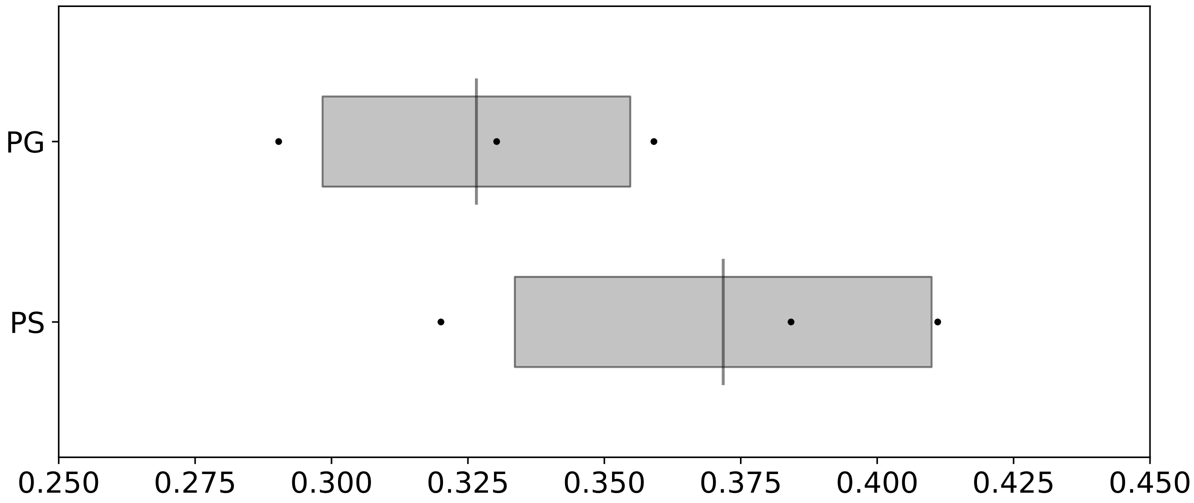


**Figure S7**: Fitness comparison between the negatively charged lipids PS and PG. The black dots are the fitness values for Membrane 3 with all PS head groups replaced by PG, referred to as PG, and Membrane 3 with all PG head groups replaced by PS, referred to as PS. The grey box marks the standard deviation, the grey line within it the mean value of the fitness.

| AMP | $\frac{\Delta x}{\text{nm}}$ | $\frac{x}{\text{nm}}$ | $\frac{x_{\text{Host}}}{\text{nm}}$ | non AMP | $\frac{\Delta x}{\text{nm}}$ | $\frac{x}{\text{nm}}$ | $\frac{x_{\text{Host}}}{\text{nm}}$ |
|---|---|---|---|---|---|---|---|
| GIHDILKYGKPS | *-0.005* | 0.914 | 0.909 | EYLIEVRESAKMTQ | -0.073 | 1.080 | 1.007 |
| GLFDVIKKVASVIGGL | 0.023 | 0.672* | 0.695$^{\dagger}$ | HILRMRIRQGMMT | -0.008 | 0.860 | 0.853 |
| FCTMIPIPRCY | 0.029 | 0.781 | 0.810 | HILRMRIRQMMT | -0.002 | 0.835 | 0.833 |
| LLPIVGNLLKSLL | 0.032 | 0.656* | 0.689$^{\dagger}$ | HIRLMRIRQMMT | 0.001 | 0.938 | 0.939 |
| FLPLILRKIVTAL | 0.045 | 0.662* | 0.707 | YQLLRIMRINIA | 0.010 | 0.794 | 0.804 |
| FLPLIGRVLSGIL | 0.047 | 0.625* | 0.672$^{\dagger}$ | GLITMLKVGLAKVQ | 0.011 | 0.753 | 0.764 |
| FVQWFSKFLGRIL | 0.051 | 0.659* | 0.710 | LRPAFKVSK | 0.012 | 0.753 | 0.765 |
| FLLFPLMCKIQGKC | 0.055 | 0.687* | 0.743 | HIRAMRIRAQMMT | 0.012 | 0.940 | 0.952 |
| GLLGPLLKIAAKVGSNLL | 0.069 | 0.558* | 0.627$^{\dagger}$ | KTLAQLSAGVKRWH | 0.015 | 0.855 | 0.869 |
| FFPIGVFCKIFKTC | 0.108 | 0.622* | 0.730 | LIQVAPLGRLLKRR | *0.040* | 0.795 | 0.834 |

**Table S1**: Verification for 25a75i. Optimal cutoff is 0.019.

#### 5.4 Convergence of the peptide membrane distance

While the exact distance between peptide and membrane varies over time, the average distance should converge over time. By approximating the positions of membrane and peptide as Gaussians (Fig 2), we can express this convergence via the central limit theorem. To explore the sampling time needed, for the Gaussians to converge, we took a look at a few example systems S8, which have been sampled for different times.

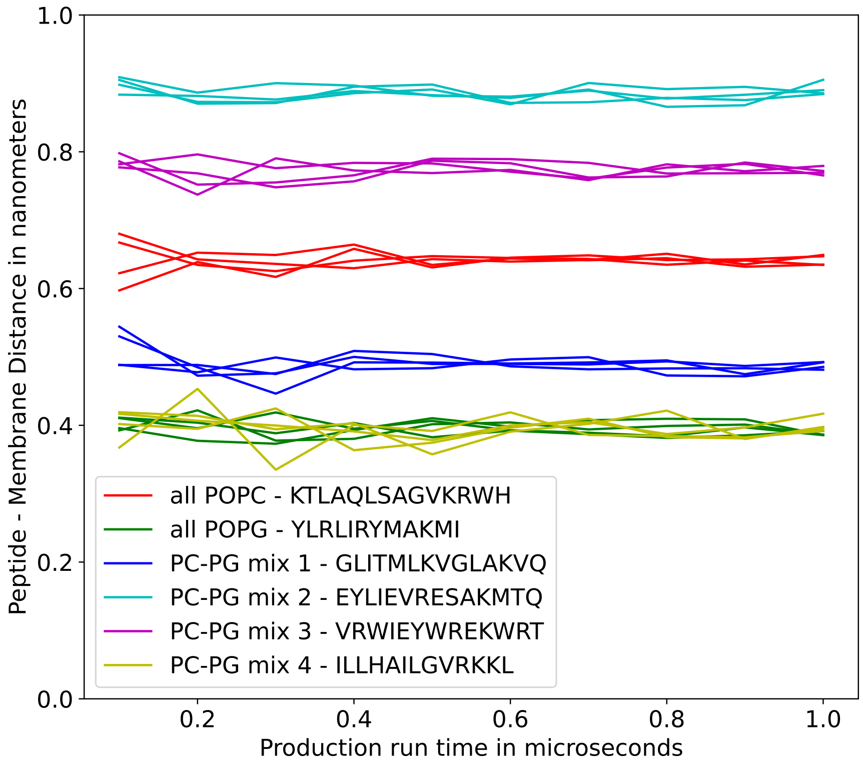


**Figure S8**: Runtime convergence check of the peptide membrane distance for reruns of different peptide membrane systems. Each system was run four times and the sampling times were evaluated between 100 ns and 1 µs in 100 nsincrements. The insertion depth appears to be converged at roughly 600 ns for all the systems. To account for potential outliers, we sampled for 900 ns in all simulations performed in this study.

| AMP | $\frac{\Delta x}{\text{nm}}$ | $\frac{x}{\text{nm}}$ | $\frac{x_{\text{Host}}}{\text{nm}}$ | non AMP | $\frac{\Delta x}{\text{nm}}$ | $\frac{x}{\text{nm}}$ | $\frac{x_{\text{Host}}}{\text{nm}}$ |
|---|---|---|---|---|---|---|---|
| ILLHAILGVRKKL | *0.009* | 0.742 | 0.752 | HRALMRIRQCMT | 0.009 | 0.849 | 0.858 |
| FPLTWLKWWKWKK | 0.029 | 0.909 | 0.939 | YRAAMLRRQYMMT | 0.020 | 0.950 | 0.969 |
| YQLRLIMKYAI | 0.047 | 0.715 | 0.762 | HRAIMLRIRQMMT | 0.022 | 0.881 | 0.904 |
| YLRLIRYMAKMI | 0.067 | 0.756 | 0.823 | VRWIEYWREKWRT | 0.023 | 0.919 | 0.941 |

**Table S2**: Rerun of the training data for 25a75i. Optimal cutoff is 0.026.

| AMP | $\frac{\Delta x}{\text{nm}}$ | $\frac{x}{\text{nm}}$ | $\frac{x_{\text{Host}}}{\text{nm}}$ | non AMP | $\frac{\Delta x}{\text{nm}}$ | $\frac{x}{\text{nm}}$ | $\frac{x_{\text{Host}}}{\text{nm}}$ |
|---|---|---|---|---|---|---|---|
| GIHDILKYGKPS | *0.036* | 0.874 | 0.909 | EYLIEVRESAKMTQ | -0.048 | 1.055 | 1.007 |
| FCTMIPIPRCY | *0.039* | 0.771 | 0.810 | HILRMRIRQGMMT | 0.029 | 0.824 | 0.853 |
| GLFDVIKKVASVIGGL | 0.078 | 0.617* | 0.695† | HIRAMRIRAQMMT | 0.031 | 0.922 | 0.952 |
| LLPIVGNLLKSLL | 0.088 | 0.600* | 0.689† | HIRLMRIRQMMT | 0.049 | 0.890 | 0.939 |
| FLLFPLMCKIQGKC | 0.108 | 0.635* | 0.743 | LRPAFKVSK | 0.054 | 0.711 | 0.765 |
| FLPLILRKIVTAL | 0.114 | 0.593* | 0.707 | HILRMRIRQMMT | 0.054 | 0.778 | 0.833 |
| FLPLIGRVLSGIL | 0.114 | 0.558* | 0.672† | KTLAQLSAGVKRWH | 0.054 | 0.815 | 0.869 |
| FVQWFSKFLGRIL | 0.128 | 0.582* | 0.710 | YQLLRIMRINIA | 0.063 | 0.742 | 0.804 |
| GLLGPLLKIAAKVGSNLL | 0.139 | 0.488* | 0.627† | GLITMLKVGLAKVQ | 0.067 | 0.697* | 0.764 |
| FFPIGVFCKIFKTC | 0.173 | 0.558* | 0.730 | LIQVAPLGRLLKRR | *0.112* | 0.723 | 0.834 |

**Table S3**: Verification for Membrane 1. Optimal cutoff is 0.072.

| AMP | $\frac{\Delta x}{\text{nm}}$ | $\frac{x}{\text{nm}}$ | $\frac{x_{\text{Host}}}{\text{nm}}$ | non AMP | $\frac{\Delta x}{\text{nm}}$ | $\frac{x}{\text{nm}}$ | $\frac{x_{\text{Host}}}{\text{nm}}$ |
|---|---|---|---|---|---|---|---|
| FPLTWLKWWKWKK | 0.075 | 0.864 | 0.939 | YRAAMLRRQYMMT | 0.030 | 0.940 | 0.969 |
| ILLHAILGVRKKL | 0.080 | 0.672* | 0.752 | VRWIEYWREKWRT | 0.053 | 0.888 | 0.941 |
| YQLRLIMKYAI | 0.103 | 0.658* | 0.762 | HRALMRIRQCMT | 0.060 | 0.798 | 0.858 |
| YLRLIRYMAKMI | 0.129 | 0.694* | 0.823 | HRAIMLRIRQMMT | 0.070 | 0.833 | 0.904 |

**Table S4**: Rerun of the training data for Membrane 1. Optimal cutoff is 0.073.

| AMP | $\frac{\Delta x}{\text{nm}}$ | $\frac{x}{\text{nm}}$ | $\frac{x_{\text{Host}}}{\text{nm}}$ | non AMP | $\frac{\Delta x}{\text{nm}}$ | $\frac{x}{\text{nm}}$ | $\frac{x_{\text{Host}}}{\text{nm}}$ |
|---|---|---|---|---|---|---|---|
| GIHDILKYGKPS | *0.041* | 0.868 | 0.909 | EYLIEVRESAKMTQ | -0.036 | 1.043 | 1.007 |
| FCTMIPIPRCY | *0.072* | 0.738 | 0.810 | HILRMRIRQGMMT | 0.033 | 0.819 | 0.853 |
| GLFDVIKKVASVIGGL | 0.090 | 0.605* | $0.695^{\dagger}$ | HIRLMRIRQMMT | 0.052 | 0.887 | 0.939 |
| LLPIVGNLLKSLL | 0.098 | 0.590* | $0.689^{\dagger}$ | LRPAFKVSK | 0.058 | 0.707 | 0.765 |
| FLPLILRKIVTAL | 0.115 | 0.591* | 0.707 | HILRMRIRQMMT | 0.059 | 0.773 | 0.833 |
| FVQWFSKFLGRIL | 0.127 | 0.583* | 0.710 | HIRAMRIRAQMMT | 0.062 | 0.891 | 0.952 |
| FLLFPLMCKIQGKC | 0.129 | 0.614* | 0.743 | YQLLRIMRINIA | 0.062 | 0.742 | 0.804 |
| FLPLIGRVLSGIL | 0.131 | 0.541* | $0.672^{\dagger}$ | GLITMLKVGLAKVQ | 0.070 | 0.694* | 0.764 |
| GLLGPLLKIAAKVGSNLL | 0.145 | 0.482* | $0.627^{\dagger}$ | KTLAQLSAGVKRWH | 0.074 | 0.795 | 0.869 |
| FFPIGVFCKIFKTC | 0.185 | 0.545* | 0.730 | LIQVAPLGRLLKRR | *0.099* | 0.735 | 0.834 |

**Table S5**: Verification for Membrane 2. Optimal cutoff is 0.082.

| AMP | $\frac{\Delta x}{\text{nm}}$ | $\frac{x}{\text{nm}}$ | $\frac{x_{\text{Host}}}{\text{nm}}$ | non AMP | $\frac{\Delta x}{\text{nm}}$ | $\frac{x}{\text{nm}}$ | $\frac{x_{\text{Host}}}{\text{nm}}$ |
|---|---|---|---|---|---|---|---|
| FPLTWLKWWKWKK | 0.081 | 0.858 | 0.939 | YRAAMLRRQYMMT | 0.063 | 0.907 | 0.969 |
| ILLHAILGVRKKL | 0.084 | 0.667* | 0.752 | VRWIEYWREKWRT | 0.064 | 0.877 | 0.941 |
| YQLRLIMKYAI | 0.116 | 0.646* | 0.762 | HRALMRIRQCMT | 0.074 | 0.784 | 0.858 |
| YLRLIRYMAKMI | 0.138 | 0.685* | 0.823 | HRAIMLRIRQMMT | 0.079 | 0.825 | 0.904 |

**Table S6**: Verification for Membrane 2. Optimal cutoff is 0.080.

| AMP | $\frac{\Delta x}{\text{nm}}$ | $\frac{x}{\text{nm}}$ | $\frac{x_{\text{Host}}}{\text{nm}}$ | non AMP | $\frac{\Delta x}{\text{nm}}$ | $\frac{x}{\text{nm}}$ | $\frac{x_{\text{Host}}}{\text{nm}}$ |
|---|---|---|---|---|---|---|---|
| GIHDILKYGKPS | *0.032* | 0.878 | 0.909 | EYLIEVRESAKMTQ | -0.043 | 1.050 | 1.007 |
| FCTMIPIPRCY | *0.064* | 0.746 | 0.810 | HILRMRIRQGMMT | 0.034 | 0.818 | 0.853 |
| GLFDVIKKVASVIGGL | 0.088 | 0.607* | $0.695^{\dagger}$ | LRPAFKVSK | 0.051 | 0.714 | 0.765 |
| LLPIVGNLLKSLL | 0.094 | 0.594* | $0.689^{\dagger}$ | HILRMRIRQMMT | 0.052 | 0.780 | 0.833 |
| FLPLILRKIVTAL | 0.108 | 0.599* | 0.707 | HIRLMRIRQMMT | 0.054 | 0.885 | 0.939 |
| FLLFPLMCKIQGKC | 0.108 | 0.635* | 0.743 | GLITMLKVGLAKVQ | 0.063 | 0.701 | 0.764 |
| FVQWFSKFLGRIL | 0.120 | 0.590* | 0.710 | KTLAQLSAGVKRWH | 0.063 | 0.806 | 0.869 |
| FLPLIGRVLSGIL | 0.124 | 0.549* | $0.672^{\dagger}$ | YQLLRIMRINIA | 0.065 | 0.739 | 0.804 |
| GLLGPLLKIAAKVGSNLL | 0.147 | 0.480* | $0.627^{\dagger}$ | HIRAMRIRAQMMT | 0.067 | 0.885 | 0.952 |
| FFPIGVFCKIFKTC | 0.179 | 0.551* | 0.730 | LIQVAPLGRLLKRR | *0.103* | 0.731 | 0.834 |

**Table S7**: Verification for Membrane 3. Optimal cutoff is 0.07765.

| AMP | $\frac{\Delta x}{\text{nm}}$ | $\frac{x}{\text{nm}}$ | $\frac{x_{\text{Host}}}{\text{nm}}$ | non AMP | $\frac{\Delta x}{\text{nm}}$ | $\frac{x}{\text{nm}}$ | $\frac{x_{\text{Host}}}{\text{nm}}$ |
|---|---|---|---|---|---|---|---|
| ILLHAILGVRKKL | *0.069* | 0.683* | 0.752 | YRAAMLRRQYMMT | 0.054 | 0.915 | 0.969 |
| FPLTWLKWWKWKK | 0.076 | 0.862 | 0.939 | VRWIEYWREKWRT | 0.062 | 0.879 | 0.941 |
| YQLRLIMKYAI | 0.111 | 0.651* | 0.762 | HRALMRIRQCMT | 0.067 | 0.791 | 0.858 |
| YLRLIRYMAKMI | 0.126 | 0.696* | 0.823 | HRAIMLRIRQMMT | 0.074 | 0.830 | 0.904 |

**Table S8**: Rerun of training data for Membrane 3. Optimal cutoff is 0.075.

| AMP | $\frac{\Delta x}{\text{nm}}$ | $\frac{x}{\text{nm}}$ | $\frac{x_{\text{Host}}}{\text{nm}}$ | non AMP | $\frac{\Delta x}{\text{nm}}$ | $\frac{x}{\text{nm}}$ | $\frac{x_{\text{Host}}}{\text{nm}}$ |
|---|---|---|---|---|---|---|---|
| FCTMIPIPRCY | *0.043* | 0.768 | 0.810 | EYLIEVRESAKMTQ | -0.037 | 1.044 | 1.007 |
| GIHDILKYGKPS | *0.045* | 0.865 | 0.909 | HILRMRIRQGMMT | 0.033 | 0.819 | 0.853 |
| GLFDVIKKVASVIGGL | 0.080 | 0.615* | $0.695^{\dagger}$ | HIRLMRIRQMMT | 0.048 | 0.891 | 0.939 |
| LLPIVGNLLKSLL | 0.094 | 0.595* | $0.689^{\dagger}$ | LRPAFKVSK | 0.050 | 0.715 | 0.765 |
| FLPLILRKIVTAL | 0.101 | 0.605* | 0.707 | HIRAMRIRAQMMT | 0.053 | 0.899 | 0.952 |
| FLLFPLMCKIQGKC | 0.112 | 0.631* | 0.743 | GLITMLKVGLAKVQ | 0.060 | 0.704 | 0.764 |
| FLPLIGRVLSGIL | 0.123 | 0.549* | $0.672^{\dagger}$ | KTLAQLSAGVKRWH | 0.065 | 0.805 | 0.869 |
| FVQWFSKFLGRIL | 0.125 | 0.585* | 0.710 | HILRMRIRQMMT | 0.065 | 0.768 | 0.833 |
| GLLGPLLKIAAKVGSNLL | 0.144 | 0.483* | $0.627^{\dagger}$ | YQLLRIMRINIA | 0.067 | 0.737 | 0.804 |
| FFPIGVFCKIFKTC | 0.164 | 0.567* | 0.730 | LIQVAPLGRLLKRR | 0.079 | 0.756 | 0.834 |

**Table S9**: Verification for Membrane 4. Optimal cutoff is 0.079.

| AMP | $\frac{\Delta x}{\text{nm}}$ | $\frac{x}{\text{nm}}$ | $\frac{x_{\text{Host}}}{\text{nm}}$ | non AMP | $\frac{\Delta x}{\text{nm}}$ | $\frac{x}{\text{nm}}$ | $\frac{x_{\text{Host}}}{\text{nm}}$ |
|---|---|---|---|---|---|---|---|
| FPLTWLKWWKWKK | 0.074 | 0.865 | 0.939 | YRAAMLRRQYMMT | 0.047 | 0.922 | 0.969 |
| ILLHAILGVRKKL | 0.076 | 0.675* | 0.752 | VRWIEYWREKWRT | 0.057 | 0.885 | 0.941 |
| YQLRLIMKYAI | 0.090 | 0.671* | 0.762 | HRALMRIRQCMT | 0.060 | 0.798 | 0.858 |
| YLRLIRYMAKMI | 0.119 | 0.703 | 0.823 | HRAIMLRIRQMMT | 0.069 | 0.835 | 0.904 |

**Table S10**: Rerun of training data for Membrane 4. Optimal cutoff is 0.071.

| AMP | $\frac{\Delta x}{\text{nm}}$ | $\frac{x}{\text{nm}}$ | $\frac{x_{\text{Host}}}{\text{nm}}$ | non AMP | $\frac{\Delta x}{\text{nm}}$ | $\frac{x}{\text{nm}}$ | $\frac{x_{\text{Host}}}{\text{nm}}$ |
|---|---|---|---|---|---|---|---|
| GIHDILKYGKPS | *0.023* | 0.887 | 0.909 | EYLIEVRESAKMTQ | -0.047 | 1.055 | 1.007 |
| FCTMIPIPRCY | *0.026* | 0.784 | 0.810 | HILRMRIRQGMMT | 0.004 | 0.848 | 0.853 |
| GLFDVIKKVASVIGGL | *0.060* | 0.635* | 0.695$^{\dagger}$ | LRPAFKVSK | 0.024 | 0.741 | 0.765 |
| LLPIVGNLLKSLL | 0.069 | 0.620* | 0.689$^{\dagger}$ | HIRAMRIRAQMMT | 0.031 | 0.921 | 0.952 |
| FLLFPLMCKIQGKC | 0.081 | 0.662* | 0.743 | HILRMRIRQMMT | 0.035 | 0.798 | 0.833 |
| FLPLIGRVLSGIL | 0.094 | 0.578* | 0.672$^{\dagger}$ | HIRLMRIRQMMT | 0.038 | 0.901 | 0.939 |
| FLPLILRKIVTAL | 0.095 | 0.612* | 0.707 | GLITMLKVGLAKVQ | 0.043 | 0.721 | 0.764 |
| FVQWFSKFLGRIL | 0.102 | 0.608* | 0.710 | YQLLRIMRINIA | 0.045 | 0.759 | 0.804 |
| GLLGPLLKIAAKVGSNLL | 0.106 | 0.521* | 0.627$^{\dagger}$ | KTLAQLSAGVKRWH | 0.048 | 0.821 | 0.869 |
| FFPIGVFCKIFKTC | 0.142 | 0.588* | 0.730 | LIQVAPLGRLLKRR | 0.060 | 0.774 | 0.834 |

**Table S11**: Verification for Membrane 5. Optimal cutoff is 0.064.

| AMP | $\frac{\Delta x}{\text{nm}}$ | $\frac{x}{\text{nm}}$ | $\frac{x_{\text{Host}}}{\text{nm}}$ | non AMP | $\frac{\Delta x}{\text{nm}}$ | $\frac{x}{\text{nm}}$ | $\frac{x_{\text{Host}}}{\text{nm}}$ |
|---|---|---|---|---|---|---|---|
| ILLHAILGVRKKL | 0.051 | 0.700 | 0.752 | YRAAMLRRQYMMT | 0.023 | 0.946 | 0.969 |
| FPLTWLKWWKWKK | 0.058 | 0.880 | 0.939 | HRALMRIRQCMT | 0.037 | 0.821 | 0.858 |
| YQLRLIMKYAI | 0.090 | 0.671* | 0.762 | VRWIEYWREKWRT | 0.040 | 0.901 | 0.941 |
| YLRLIRYMAKMI | 0.107 | 0.716 | 0.823 | HRAIMLRIRQMMT | 0.048 | 0.856 | 0.904 |

**Table S12**: Rerun of training data for Membrane 5. Optimal cutoff is 0.049.
